# Not all DNase 1 are created equal

**DOI:** 10.64898/2026.11.23.768681

**Authors:** Deepali Lehri, Janice B Fung, Madison A Spears, Daniel G Kopulos, Ravi Sachidanandam

**Affiliations:** New York Medical College, Valhalla, NY 10595

## Abstract

DNase I is a ubiquitous reagent, used to remove contaminating DNA from RNA preparations and other samples, and preparations from different suppliers are widely treated as interchangeable. Suppliers describe their products as improved — faster, more complete, more stable — but what *better* means is rarely defined in terms a user can reproduce: activity units are tied to each supplier’s own assay, recommended protocols specify a single amount and a single time, and stability in storage, through freeze–thaw cycles and at working temperature, is seldom quantified.

Here we adapt a low-cost fluorophore–quencher reporter of the kind used for Cas12 trans-cleavage diagnostics as a kinetic assay for DNase I, and compare preparations from three suppliers, each used at 1 U/µL, on plates that carry blanks, replicates and a standard of fully digested reporter. At large amounts (16– 32 U) ThermoFisher and Zymo preparations were indistinguishable early in the reaction. At sub-saturating amounts (≤ 4 U) the product formed per labelled unit differed up to ≈ 3.6-fold at 1 U (Zymo over ThermoFisher) and by considerably more at 0.25–0.5 U, where ThermoFisher signal was close to background. The ranking depended on dose, on time, and on whether rate or yield was the metric: from about 3 h onward ThermoFisher formed the least product at 1–4 U yet had the shortest fitted time constants, Synthego and Zymo were similar at ≤1 U, and Synthego, slower to start, yielded more than Zymo after about 2–3 h at 4 U. On a single plate, two Zymo DNase I aliquots from different laboratories differed up to 19-fold at equal nominal volume.

We argue that DNase I performance should be reported as calibrated dose–response and time-course data under stated conditions, with storage stability quantified, and that laboratories should verify the enzyme they actually hold. The reporter costs about $0.10 per reaction, is in principle readable on a simple LED fluorometer, and can be modified to probe dsDNA, RNA and nucleic-acid–binding proteins.

## 2. Introduction

Deoxyribonuclease I (DNase I) is a divalent-cation-dependent endonuclease that hydrolyzes the phospho-diester backbone of DNA [1, 2]; in the presence of Mg^2+^ it nicks each strand of double-stranded DNA independently, generating random cleavage sites [3]. Beyond its routine use for removing contaminating DNA during RNA purification and other sample preparation [3, 4], DNase I activity underlies apoptotic DNA fragmentation and autoimmune regulation of extracellular DNA [5], and genome-wide mapping of regulatory DNA via DNase I hypersensitivity [6]. It is sold by essentially every major life-science supplier — Zymo, Thermo Fisher, NEB, Synthego, and others — and is commonly treated as an interchangeable commodity reagent.

Preparations differ in origin, formulation and, in some cases, engineering, and suppliers describe them as improved: faster, more complete in digestion, or more stable. What “better” means, however, is rarely stated in terms a user can reproduce. An activity unit is defined by a supplier’s own assay (substrate, buffer, temperature and endpoint), and such definitions have historically differed [7]. Recommended protocols give a single amount and a single incubation time — for example 2 units for 10 minutes at 37°C [4] — generally chosen to ensure complete digestion, the regime in which differences between preparations are hardest to see. And stability, meaning how long a preparation keeps its activity in storage, through freeze–thaw cycles and at working temperature, is to our knowledge seldom quantified in a form a user can act on, although DNase I is known to lose activity at physiological temperature [8]. Two tubes carrying the same label are therefore treated as equivalent, and a laboratory cannot tell, without measuring it, whether the tube in its freezer behaves like the one its protocol was written for.

Many assays for DNase I activity exist: DNA–methyl green and hyperchromicity methods [7], fluorophore– quencher oligonucleotide assays for high-throughput inhibitor screening [9], graphene-oxide, gold-nanoparticle and DNAzyme colorimetric platforms [10, 11, 12, 13], and electrochemical biosensors [14]. They have largely been developed for drug screening and clinical diagnostics, in tumor tissue [10], serum [12] and saliva [14], and have not, to our knowledge, been used to compare commercial preparations side by side across amounts and times.

A FRET-based Cas12 single-stranded DNA (ssDNA) reporter is a short oligonucleotide labeled with a fluorescent donor and a quencher (we use standard pairs such as FAM and BHQ). **Collateral cleavage**: when the Cas12-crRNA complex binds its specific target DNA, it activates non-specific trans-cleavage of nearby single-stranded DNA [15, 16]. **Signal generation**: the activated Cas12 cuts the reporter, separating the fluorophore from the quencher [17]. **Readout**: fluorescence is restored in proportion to the amount of cleavage [15] (Fig. 1A). Our own work began with Cas12 detection of double-stranded DNA (dsDNA) targets, confirmed with a gBlock substrate resolved on a TapeStation. We then wanted to test activated Cas12’s collateral activity on ssDNA with this reporter, and needed a positive control for reporter cleavage independent of Cas12. DNase I, an orthogonal enzyme known to cleave ssDNA, was a natural choice. Characterizing DNase I from several sources in our institute in 96-well plates (Fig. 1B) showed that stocks of nominally the same activity produced very different amounts of signal, which led to the calibrated comparison reported here.

**Figure 1:**
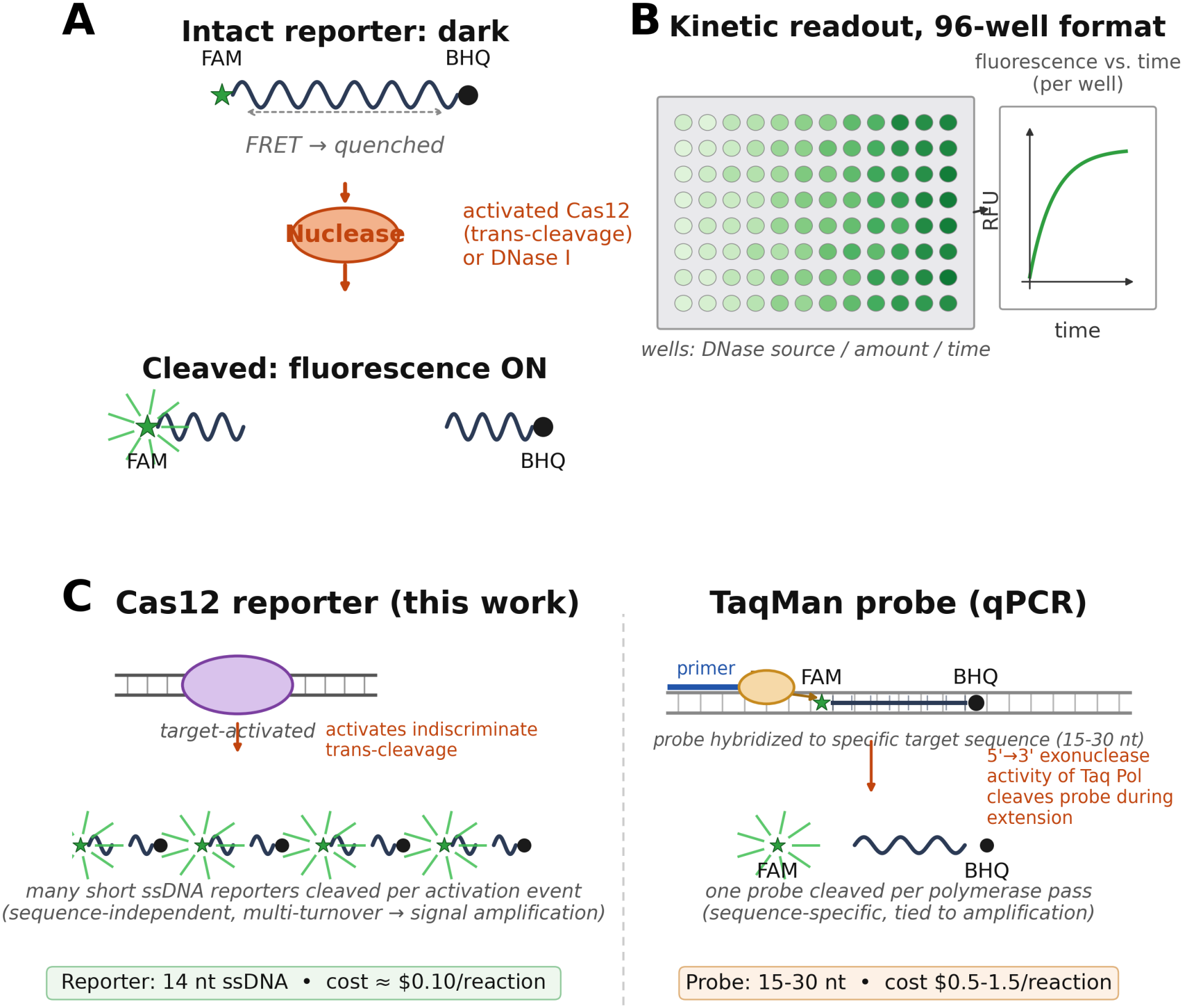
The Cas12/DNase reporter assay. (**A**) Reporter design and readout: an intact ssDNA oligo bridging a fluorophore (FAM) and quencher (BHQ) is dark via FRET; nuclease cleavage (Cas12 collateral trans-cleavage, or DNase I as used throughout this paper) separates the pair and restores fluorescence. (**B**) The assay is run in 96-well plates, with fluorescence read repeatedly over time per well to yield the kinetic curves fit throughout this paper. (**C**) Mechanistic comparison with the TaqMan probe used in the most directly comparable prior work [18]: both use a fluorophore-quencher oligo, but a Cas12 reporter is short and sequence-independent, cleaved indiscriminately and repeatedly by a single activated Cas12 (multi-turnover signal amplification), whereas a TaqMan probe is long, sequence-specific, and cleaved once per polymerase pass via the 5’→3’ exonuclease activity of Taq polymerase during PCR extension – a difference reflected in per-reaction cost (Table 1).

The assay most directly comparable to ours uses a TaqMan fluorophore–quencher probe to measure sputum DNase I kinetics in asthma, where reduced airway DNase activity was linked to more severe disease and mucus plugging, in part via carbamylation of the enzyme [18]. TaqMan probes use the same basic readout, but differ in sequence composition, in the mechanism that cleaves them, and in cost (Fig. 1C, Table 1). Being long, sequence-specific reagents, they are not efficient Cas12 substrates, and the two chemistries are not interchangeable. Despite the widespread use of Cas12 reporters in diagnostics such as DETECTR, we are not aware of prior work using one to quantify DNase I kinetics.

**Table 1:** Cas12 reporter vs TaqMan probe.

| **Feature** | **Cas12 reporter** | **TaqMan probe** |
|---|---|---|
| Structure | Fluorophore and quencher on a short ssDNA (14 nt here) | Fluorophore and quencher on a 15–30 nt oligonucleotide complementary to an internal region of the PCR product |
| Specificity | Sequence-independent; cleaved indiscriminately once Cas12 is activated | Sequence-dependent; must bind the target amplicon to be cleaved |
| Cleavage | Trans-cleavage (collateral) activity of Cas12 | 5′ →3′ exonuclease activity of Taq polymerase during extension |
| Primary use | CRISPR-based diagnostics (e.g., DETECTR) | Real-time quantitative PCR |
| Probe cost | Generic sequence; about \$280 for ≈3000 reactions in our hands | Custom; \$120–\$400 per probe (5–20 nmol) |
| Cost per reaction | ≈\$0.10 (reporter only) | ≈\$0.5–\$1.5 |

Here we use this reporter to compare DNase I from three suppliers, each used at 1 U/µL, at amounts from 0.25 to 32 units, and ask what a user would need to know to compare them: how much product forms, how fast, and for how long the enzyme stays active. We do not measure storage stability here; our aim is to show what a cheap assay can report.

## 3. Materials and Methods

### 3.1 Reporter assay and plate layout

DNase activity was measured with a ssDNA FRET reporter bearing FAM and BHQ connected by a short ssDNA linker (/56-FAM/TTATTTATTTATTT/3BHQ 1/, from IDT); cleavage of the linker increases fluorescence. Plates (96-well) were read on a BioTek Synergy HT reader (485/20 nm excitation, 528/20 nm emission, top optics, gain 35, read height 4.5 mm), as end-point reads of the full plate repeated over time. In Exp3 and Exp4 each plate carried DNase I wells in replicate rows, no-enzyme controls (reporter only), blank wells (no reporter, no enzyme), and wells containing the reporter after complete digestion of its linker at the working concentration (“fluor1”) and at half that (“fluor0p5”), used as an on-plate calibration standard. Exp1 and Exp2 were earlier single-well pilots without blanks or standards (Table 2). The reporter concentration was µM unless stated (Table 2).

**Table 2:** Experiments. Reads: number of plate reads; last read: hours after the first read. Doses are labelled Units (= µL at 1 U/µL).

| **Exp** | **Plate date** | **DNase I (doses, U)** | **Reporter (µM)** | **Wells/cond.** | **Blanks, standard** | **Reads** | **Last read** |
|---|---|---|---|---|---|---|---|
| Exp1 | 5 Aug 2026 | T (2.5, 5, 10); W (10) | 0.83–5 | 1 | no | 11 | 24.1 h |
| Exp2 | 19 Aug 2026 | T (10); W, G (1.25, 2.5, 5) | 5 | 1 | no | 11 | 24.8 h |
| Exp3 | 10 Sep 2026 | T (0.25–16); Z (0.625–32) | 2.5 (T 4 U, Z 4 and 16 U also 1.25) | 3 | yes | 8 | 23.0 h |
| Exp4 | 17 Sep 2026 | T, Z (0.25–0.75); S (0.25–4) | 2.5 | 1–3 | yes | 9 | 21.9 h |

### 3.2 DNase I preparations and units

T is the laboratory’s own ThermoFisher DNase I stock, stored at − 20°C and never freeze–thawed; Z and S are freshly received Zymo and Synthego stocks. S is supplied at 4 U/µL and was diluted with nuclease-free water to 1 U/µL before use; T and Z are labelled 1 U/µL. W and G are Zymo DNase I aliquots obtained through two other laboratories (1 U/µL assumed). Amounts of 0.25–32 µL were added per well, so labelled Units equal microlitres. Labelled Units are used for comparison because they are what a user has; they are defined by each supplier’s own assay and are not assumed to be equivalent on our substrate. For each DNase we used the buffer provided by the manufacturer.

### 3.3 Reaction setup and read times

Reporter and enzyme were combined in the assay plate, and the plate was read repeatedly. In Exp3 and Exp4 enzyme was dispensed with a multichannel pipette, one plate column at a time; in Exp1 and Exp2 well by well. Read times are the date and time stamps recorded in each plate-reader file, in minutes since the first read of that plate (Table ST1). The first read is not time zero of the reaction; because the model below has a free baseline, a constant offset is absorbed in the fitted baseline and amplitude and does not bias the time constant.

### 3.4 Signal normalization

Blank-well signal (mean of all blanks at the same read) was subtracted in Exp3 and Exp4. Signal was then expressed as a percentage of the fully digested standard “fluor1” on the same plate at the same read (mean of 4 wells in Exp3, 2 in Exp4), an index of the fraction of reporter cleaved. The standard was run at two levels and was not linear: half concentration (“fluor0p5”) gave 38% and 39% of the full signal on the two plates, so the index understates the fraction cleaved where signal is low, and we report it as a calibrated index rather than convert it to amounts. Wells with 1.25 µM reporter were scaled to “fluor0p5”.

### 3.5 Kinetic model and uncertainty

Each well’s time course was fit to *F* (*t*) = *F*_0_+*A* (1 −*e*^−*t/τ*^) (rate constant *k* = 1*/τ*) by bounded nonlinear least squares. The 95% interval for *τ* was obtained by profiling the residual sum of squares over *τ* (2–50,000 min) and applying the *F* -test limit for one parameter. A fit passed quality control if *R*^2^ ≥ 0.9, *A* ≥ 3 RFU and *F*_0_ was not at a bound; *τ* was called *reliable* if the fit passed and its interval spanned no more than a factor of three without reaching the end of the grid. Rate constants are quoted only for reliable fits. The model summarizes a sparsely sampled curve and is not a mechanism; where the reaction is largely complete by the first read, or the signal is near background, *τ* is not determined.

### 3.6 Replicates, statistics and software

Exp3 and Exp4 used replicate wells (Table 2); signal is the mean of wells and *τ* the mean of reliable wells with the mean 95% interval. Exp1 and Exp2 had one well per condition and are treated as pilot data. Parsing, analysis and figures used Python (numpy, scipy, pandas, matplotlib). Plate-reader files, layouts, and the tidy time-series and fit tables are provided as supplementary data (Data availability).

## 4. Results

### 4.1 Plate calibration

The fully digested standard read 13 ± 2 5 RFU (fluor1) and 51 ± 2 RFU (fluor0p5) on the Exp3 plate and 55 ± 6 and 21.5 ± 1.4 RFU on the Exp4 plate, a 2.4-fold difference in sensitivity at both levels. On both plates half the concentration gave 38–39% of the full signal rather than 50%. Expressing signal as a percentage of each plate’s own standard reconciled the plates: fresh Zymo at 0.5–0.75 U on Exp4 gave 16.7% and 18.6% at 3 h and 15.8% and 17.6% at 22 h, against 14.4% and 17.8% at 0.625 U on Exp3, and ThermoFisher at 0.25 U gave 1.4% and 1.6% at 22 h on the two plates. Wells with 1.25 µM reporter, scaled to their own standard, agreed with the matching 2.5 µM wells within 7–20% at 3–22 h. Plates are therefore compared through the standard-normalized signal, at the reads that occurred at matched actual times (≈24 min, ≈3 h and ≈22 h; Table ST1).

### 4.2 Performance at matched amounts depends on dose, time and source

With all three preparations used at 1 U/µL, the product formed per labelled unit was not the same (Fig. 2, Table 3). Values below are percentages of the fully digested standard.

**Table 3:** Calibrated signal per labelled unit. Blank-subtracted signal as a percentage of the on-plate fully digested reporter standard (mean of wells), at actual times ≈0.4 h, ≈3 h and ≈22 h after the first read. T, ThermoFisher; Z, fresh Zymo; S, Synthego. Entries come from plate Exp3 (T ≥1 U, Z ≥0.625 U), plate Exp4 (S; T and Z ≤0.75 U) or both (T at 0.25 U); “–” not measured. Reporter 2.5 µM.

|  | ≈0.4 h |  |  | ≈3 h |  |  | ≈22 h |  |  |
|---|---|---|---|---|---|---|---|---|---|
| Units | T | Z | S | T | Z | S | T | Z | S |
| 0.25 | 1.6 | 5.1 | 1.4 | 1.5 | 12.9 | 11.0 | 1.5 | 11.4 | 11.7 |
| 0.5 | 2.0 | 6.0 | 4.5 | 3.3 | 16.7 | 14.1 | 2.8 | 15.8 | 14.0 |
| 0.625 | – | 5.3 | – | – | 14.4 | – | – | 17.8 | – |
| 0.75 | 3.2 | 5.1 | 7.6 | 9.7 | 18.6 | 17.9 | 5.8 | 17.6 | 17.0 |
| 1 | 2.5 | 7.1 | 6.4 | 6.1 | 20.8 | 20.5 | 7.6 | 27.0 | 21.1 |
| 2 | 6.3 | – | 8.2 | 17.0 | – | 28.7 | 18.7 | – | 32.3 |
| 4 | 18.8 | 19.8 | 12.6 | 41.3 | 55.8 | 61.7 | 42.9 | 60.0 | 73.0 |
| 8 | 40.9 | – | – | 61.9 | – | – | 62.2 | – | – |
| 16 | 65.4 | 66.9 | – | 73.5 | 87.2 | – | 73.5 | 84.0 | – |
| 32 | – | 94.6 | – | – | 90.5 | – | – | 82.8 | – |

**Figure 2:**
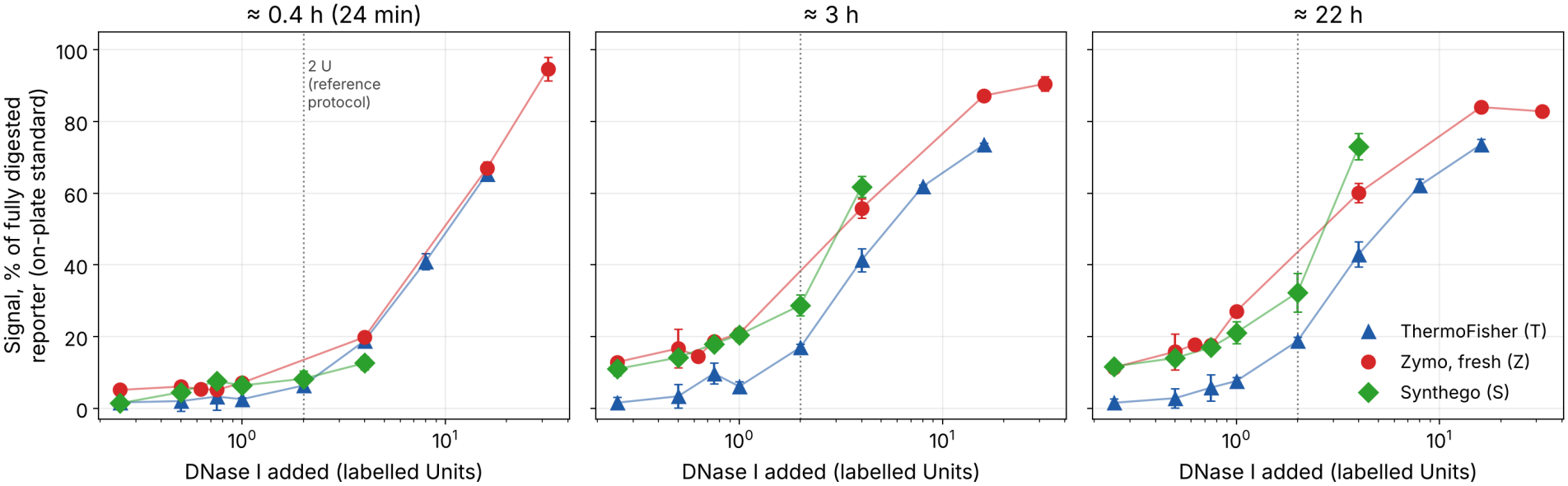
Performance per labelled unit depends on dose, time and source. Signal (blank-subtracted, as a percentage of the fully digested reporter standard on the same plate at the same read; mean *±* SD of replicate wells) against DNase I amount (all stocks at 1 U/µL), at matched actual read times of ≈24 min, ≈3 h and ≈22 h after the first read. Dotted line, 2 U (amount in a reference protocol [4]). Reporter 2.5 µM.

#### At large amounts the preparations converge

At 16 U, ThermoFisher and Zymo gave 65% and 67% by ≈ 24 min, and 74% and 84–87% by 3–22 h; Zymo at 32 U reached 95% by 24 min. Early in the reaction the preparations were indistinguishable at high amounts, which is where single-dose protocols are likely to operate.

#### At sub-saturating amounts the differences are large

At 1 U on Exp3, Zymo formed 2.8, 3.4 and 3.6 times as much signal as ThermoFisher at ≈24 min, 3 h and 22 h (7.1%, 20.8%, 27.0% against 2.5%, 6.1%, 7.6%). At 0.25–0.75 U ThermoFisher produced ≤ 10% of the standard even after 22 h, against 11–19% for Zymo and Synthego. At 4 U ThermoFisher reached 43% by 22 h against 60% for Zymo and 73% for Synthego.

#### Synthego and Zymo behaved alike at low amounts and differed in time course at 4 U

From 0.25 to 1 U their signals at 3 h agreed within 16% (for example 14.1% and 16.7% at 0.5 U, 20.5% and 20.8% at 1 U), and their time courses at 1 U were nearly superimposable (Fig. 3). At 4 U Synthego started more slowly (12.6% against 19.8% for Zymo and 18.8% for ThermoFisher at 24 min) but overtook both after about 2–3 h (61.7%, 55.8% and 41.3% at 3 h; 73.0%, 60.0% and 42.9% at 22 h).

**Figure 3:**
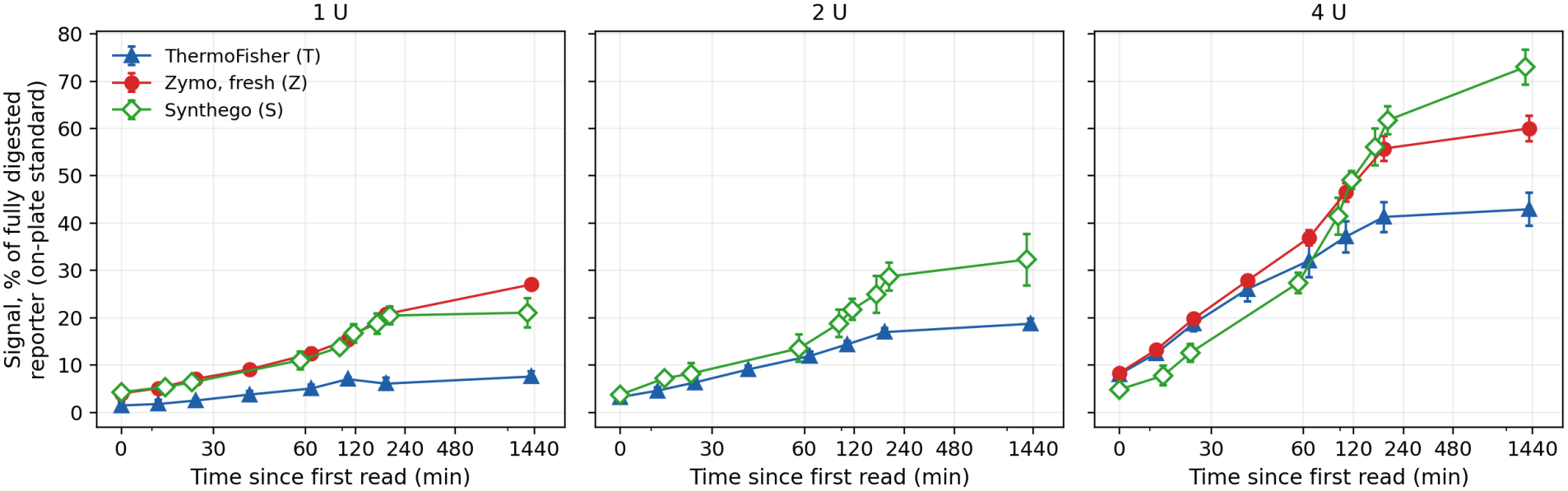
Time courses at 1, 2 and 4 U. Standard-normalized signal against actual time since the first read (axis linear to 60 min, logarithmic beyond; mean *±* SD of replicate wells). At 1 U the Zymo and Synthego curves are nearly superimposable and ThermoFisher is about threefold lower; at 2 U Synthego exceeds ThermoFisher after the first reads; at 4 U the three are within about 1.6-fold of one another during the first hour, after which ThermoFisher plateaus lowest and Synthego highest. The last point on each curve was read the next day.

*At 2 U, the amount in a widely used reference protocol [4]*, Synthego formed more signal than ThermoFisher at every read (1.3 ×, 1.7× and 1.7× at 24 min, 3 h and 22 h; 8.2%, 28.7%, 32.3% against 6.3%, 17.0%, 18.7%); Zymo was not measured at 2 U, and its values at 1 and 4 U bracket that amount. All preparations were far from complete at 2 U: after 22 h at most 19–32% of the standard had been formed. Our reaction conditions differ from that protocol’s, so these numbers do not predict how completely a particular RNA preparation would be cleared, only that the preparations differ at that amount.

### 4.3 Time constants

From 4 U upward, *τ* shortened with dose for both ThermoFisher and Zymo (Fig. 4, Table ST2; *τ* = 66→ 36 → 20 min at 4, 8 and 16 U for ThermoFisher; 93 → 24 min at 4 and 16 U for Zymo), and at 32 U Zymo had finished before the first read. At and below about 1 U, *τ* was long (≈ 90–170 min) and nearly independent of dose, with overlapping intervals: 93–116 min for Zymo at 0.25–0.75 U and 162–165 min at 0.625–1 U, and 91–139 min for Synthego across 0.25–4 U. The signal at long times also rose less than in proportion to dose in this range (for Synthego 11.7% at 0.25 U against 21.1% at 1 U at 22 h, a 1.8-fold rise for a four-fold increase in amount).

**Figure 4:**
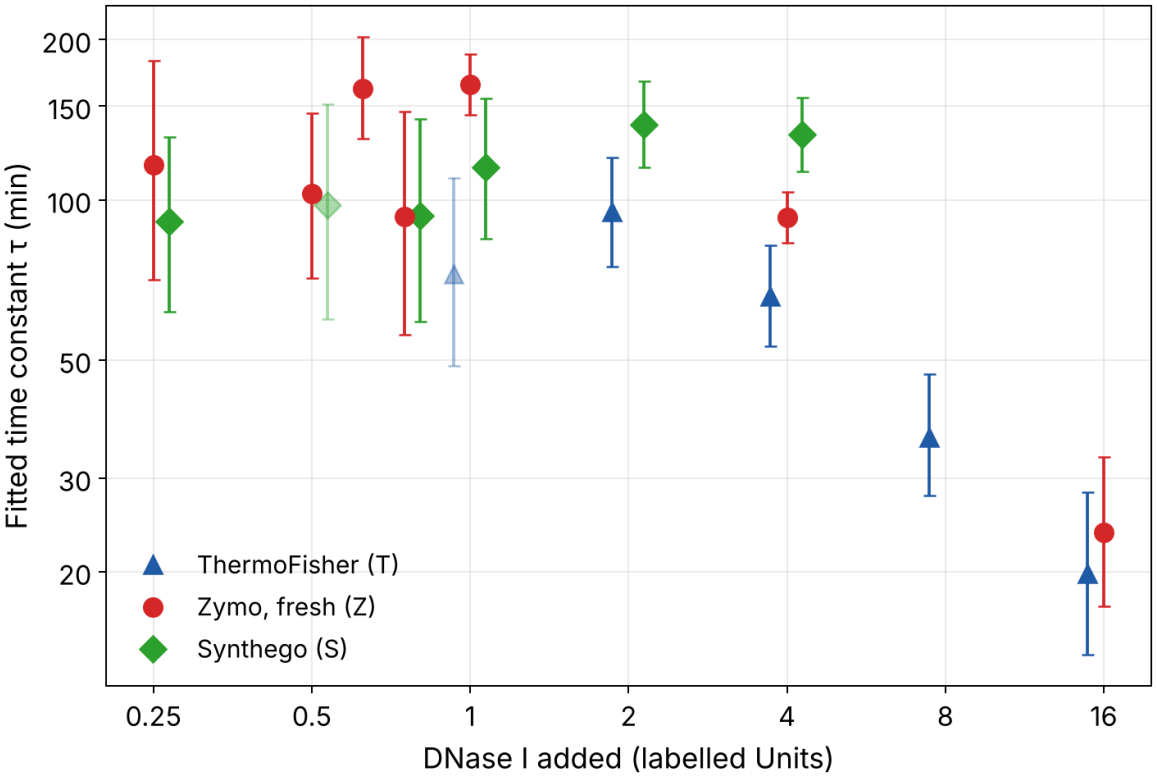
Fitted time constants against amount. Mean fitted *τ* of wells with a reliable fit at each labelled amount, with the mean 95% profile interval. Faded symbols: fewer than half of the wells in the condition had a reliable fit. ThermoFisher below 1 U and Zymo at 32 U are omitted (no passing fit).

Rate and yield gave different rankings. By rate constant ThermoFisher was equal to or faster than Zymo at every matched amount (*k* = 0.92 against 0.65/h at 4 U, 3.03 against 2.53/h at 16 U), yet it formed less product at every matched amount from 1 to 16 U by 22 h. At the small amounts less than about 30% of the reporter had been cleaved when the signal levelled off, so exhaustion of substrate does not end the reaction there. We therefore use fixed-time yield on the calibrated scale as the primary comparison, and quote *τ* only where its interval is bounded.

### 4.4 Aliquots of undocumented history, and run-to-run variation

The pilot experiments (single wells, no blanks or standards; Table ST3) are used only for comparisons within a plate. On Exp2, the Zymo aliquot from one laboratory (G) formed far less signal than the Zymo aliquot from another (W) at the same nominal volume: at ≈ 134 min, 19, 11 and 2.6 times less at 1.25, 2.5 and 5 µL (6, 13 and 47 RFU against 113, 139 and 121 RFU), and 13.5, 6.6 and 1.6 times less at 25 h. The storage history and concentration after any dilution of these aliquots are not documented, so the difference cannot be assigned to a cause.

The same ThermoFisher stock, measured on two plates five weeks apart, gave similar product but different kinetics. The fitted amplitude on Exp1 at 2.5, 5 and 10 U (26, 45, 61 RFU) matched the Exp3 amplitudes interpolated to those amounts (29, 53, 59 RFU), but the rate constant was 1.9–2.4 times higher on Exp1 (*k* = 1.35, 2.69 and 4.25/h against 0.71, 1.11 and 2.03/h). The plates were run at different temperatures (24.7–25.7°C on Exp1 and 21.0–22.8°C on Exp3 during the first 3 h), which could account for part of this. The variation is as large as most differences between suppliers in Fig. 2.

## 5 Discussion

### 5.1 What “better” has to mean

Our data show three separable properties that a single label cannot carry: how quickly product appears (the time constant), how much appears (yield at a stated time and amount), and for how long the enzyme remains active under the conditions of use. The three preparations ranked differently on each. ThermoFisher had the shortest time constants but, from about 3 h onward, formed the least product per labelled unit at 1–4 U; Zymo and Synthego formed similar amounts at 0.25–1 U; Synthego had the longest time constants at 4 U and the highest yield after about three hours. None of this is visible at a single large amount and a long time (at 16 U early signals were 65% and 67%), which is where recommended protocols tend to operate. Claims of a “better” enzyme therefore cannot be compared from labels, and may all be true under the conditions each supplier chose. What is needed is the dose–response and time course on a stated substrate under stated conditions, with the amounts that matter for the application included. Because the ThermoFisher stock was the laboratory’s own, and of unrecorded age, its lower yield may partly reflect age rather than the product; the central point, that labelled Units and manufacturer identity did not predict performance on a common substrate, does not depend on this.

### 5.2 The low-amount regime

At sub-saturating amounts the time constant was long and nearly independent of amount, and for Synthego the yield rose little between 0.25 and 1 U. Several explanations are consistent with this: loss of enzyme activity during the reaction through inactivation or adsorption to plastic, which removes a larger fraction of a small total amount of protein; a more readily cleaved subpopulation of reporter that dominates the signal when enzyme is limiting; a fixed inhibitory component such as a chelator carried over from the storage buffer; sparse enzyme-substrate encounters in an unmixed well; or product inhibition by released fragments. True cooperativity is the least likely, since DNase I is a single ∼ 37 kDa polypeptide with one catalytic site [3]. Substrate depletion is excluded, since less than about 30% of the reporter was cleaved. These can be separated by pre-incubating enzyme in assay buffer without reporter before adding it, by spiking fresh enzyme late in the reaction, by adding carrier protein or using low-binding plastic, and by denser sampling in amount; the same experiments would give the stability information that is seldom reported.

### 5.3 What calibrated reporting would involve

A lot of DNase I should be treated as a reagent whose activity on the user’s own substrate is measured, not assumed. Our reporter costs about $0.10 per reaction (about $280 for ≈ 3000 reactions; Table 1), several-fold less than TaqMan-based alternatives, and the cost scales with the amount of reporter used per well, so routine verification is realistic. A laboratory could record a baseline for a single reference condition when a lot is received and re-run only that condition periodically, or before a demanding digestion, to detect drift.

For comparisons between preparations and laboratories we suggest: (1) report the substrate, buffer including divalent-cation concentrations, volume and temperature; (2) report a dose–response including subsaturating amounts, and a time course, not a single point; (3) put blanks, replicates and a standard of fully digested reporter on every plate, at several concentrations spanning the working range, and express signal relative to it; (4) put a reference preparation on every plate, since the same ThermoFisher stock varied about twofold in rate constant between two plates (Section 4.4); (5) use the instrument’s recorded read times or automated kinetic reads; and (6) report the identity, lot, age, storage and freeze–thaw history of the enzyme, and its concentration in U/µL as supplied and as used. Suppliers sell DNase I at different concentrations (the Synthego preparation here is supplied at 4 U/µL, the others at 1 U/µL), so a volume cannot be converted to units, or compared between products, without it. Suppliers should state concentration and unit definition prominently, state what “improved” means — speed, completeness, specific activity on a defined substrate, or stability — and under what conditions, and publish stability in the forms users need: shelf life at the storage temperature, tolerance of repeated freeze–thaw cycles, and loss of activity at working temperature.

### 5.4 Limitations

The ThermoFisher stock was the laboratory’s own, stored at − 20°C without freeze–thaw cycles but of unrecorded age, whereas Zymo and Synthego were freshly received, and the W and G aliquots had undocumented histories; the study compares preparations as held, not manufacturers as shipped. Synthego was diluted from 4 to 1 U/µL before use, a step the other preparations did not undergo. The fully digested standard was run at two concentrations and was not linear, so the conversion from signal to fraction cleaved is approximate at low signal. Plates were at different, uncontrolled temperatures (about 21–26°C), reads were sparse, and long-time points are from the next day, so *τ* is bounded only within wide intervals. ThermoFisher below 1 U and Zymo at 32 U could not be fitted, and Synthego was not tested above 4 U. The reporter is a short ssDNA oligonucleotide whereas much use of DNase I targets dsDNA, so rankings here need not hold on dsDNA. The amounts compared to the reference protocol [4] were chosen because that recipe is widely used and brand-agnostic, not because any supplier recommends it. Stability in storage was not tested.

### 5.5 Outlook: a generalizable reporter platform

Nothing in the reporter design is specific to DNase I or to an ssDNA linker: the readout depends only on an activity cleaving the oligonucleotide that bridges a fluorophore and its quencher. A dsDNA-linked version would assay the same preparations on a dsDNA substrate alongside their ssDNA kinetics on the same plate, and ssRNA and dsRNA linkers would extend the approach to RNases, which are also common reagents whose lot-to-lot activity is rarely characterized by end users. Linker cleavage could also serve as a kinetic readout of sequence-specific protein binding: a bound protein protects the linker from a reference nuclease, reducing the rate in proportion to occupancy, a continuous, plate-based analogue of DNase I footprinting [19] and RNase protection assays [20]. We have not generated data toward these extensions, and a protection design needs the reference nuclease to be characterized first, which is what this paper shows is necessary.

FRET is one way to couple cleavage to fluorescence. Single-label reporters such as DNA-templated silver nanocluster probes [21] remove the second label and its cost; a short, generic, single-label reporter could in principle combine both savings. Nor is a benchtop plate reader fundamental: an open-source LED fluorescence reader built for under $15 has been used with Cas13a trans-cleavage reporters at a limit of detection of ≈3.7– 6.8 nM [22], with the expected trade-off in sensitivity and throughput, and would allow per-lot spot checks without a plate reader.

## 6 Conclusion

DNase I from different suppliers, all used at 1 U/µL, did not behave alike on a common substrate, and the differences depended on how much enzyme was used, when it was measured, and whether rate or yield was compared. The differences were hidden at large amounts and late times, where many protocols tend to operate, and large at the sub-saturating amounts that a user might use in sensitive experiments. Two Zymo DNase I aliquots from different laboratories differed up to 19-fold on one plate, and the same ThermoFisher stock measured five weeks apart varied about twofold in rate. Labels, unit definitions and single-point protocols did not predict any of this.

We therefore recommend that DNase I performance be reported as calibrated dose–response and time-course data under stated conditions, that suppliers define what “improved” means, state concentration clearly and publish stability in storage, through freeze–thaw cycles and at working temperature. It would be in the laboratories’ best interest to measure the enzyme they actually hold, using a cheap on-plate-calibrated assay of the kind described here. The same reporter, modified for dsDNA, RNA or nucleic-acid–binding proteins, offers a route to the same kind of calibrated comparison for other enzymes.

## 7 Acknowledgements

Dr. Jan Geliebter gave us helpful comments. Claude (Anthropic) was used for help with editing the manuscript and some coding for the plots. The authors checked all work and take responsibility for the paper.

## Glossary

DNase 1: Deoxyribonuclease
FRET: Förster Resonance Energy Transfer
FAM: Fluorescein, a popular fluorescent reporter dye
BHQ: Black Hole Quencher, a popular dark quencher
ssDNA: single-stranded DNA
dsDNA: double-stranded DNA
RFU: relative fluorescence unit (plate-reader output)
U: unit of DNase I activity as labelled by the supplier

## 8 Data Availability

Plate-reader exports, plate layouts, and the tidy time-series and per-well fit tables (DNase_all_experi ments_timeseries.csv, DNase_all_experiments_fits.csv, with a data dictionary) are provided as supplementary data (https://katahdin.girihlet.com/dnase_suppl/).

## S1 Supplementary Material

**Table ST1:** Read times. Minutes since the first read of each plate, computed from the date and time stamp recorded in each plate-reader file.

| **Exp** | **Date** | **Read times since first read (min)** |
|---|---|---|
| Exp1 | 05 Aug 2026 | 0, 12, 25, 38, 50, 63, 91, 114, 142, 197, 1447 |
| Exp2 | 19 Aug 2026 | 0, 16, 28, 40, 52, 63, 88, 111, 134, 238, 1487 |
| Exp3 | 10 Sep 2026 | 0, 12, 24, 42, 65, 108, 184, 1382 |
| Exp4 | 17 Sep 2026 | 0, 14, 23, 58, 97, 117, 163, 194, 1312 |

**Table ST2:** Fits for Exp3 and Exp4 (2.5 µM reporter). Wells: wells in the condition; QC: passing quality control; rel.: wells with a reliable *τ* . *τ* : mean of reliable wells (mean 95% profile interval); *k* = 1*/τ* (mean *±* SEM of reliable wells); *A*: mean fitted amplitude of passing wells (RFU; Exp3 raw, Exp4 blank-subtracted; not comparable between plates).

| Exp | Src | U | Wells | QC | rel. | *τ* (min) | *k* (/h) | *A* |
|---|---|---|---|---|---|---|---|---|
| Exp3 | T | 0.25 | 3 | 0 | 0 | – | – | – |
| Exp3 | T | 1 | 3 | 2 | 1 | 73 (49–110) | 0.83 | 9.8 |
| Exp3 | T | 2 | 3 | 3 | 3 | 95 (75–120) | 0.63 ± 0.01 | 22.2 |
| Exp3 | T | 4 | 3 | 3 | 3 | 66 (53–82) | 0.92 ± 0.05 | 49.3 |
| Exp3 | T | 8 | 3 | 3 | 3 | 36 (28–47) | 1.68 ± 0.06 | 61.8 |
| Exp3 | T | 16 | 3 | 3 | 3 | 20 (14–28) | 3.03 ± 0.06 | 53.2 |
| Exp3 | Z | 0.625 | 3 | 3 | 3 | 162 (130–203) | 0.37 ± 0.01 | 21.4 |
| Exp3 | Z | 1 | 3 | 3 | 3 | 165 (144–188) | 0.36 ± 0.01 | 33.2 |
| Exp3 | Z | 4 | 3 | 3 | 3 | 93 (83–103) | 0.65 ± 0.01 | 74.6 |
| Exp3 | Z | 16 | 3 | 3 | 3 | 24 (17–33) | 2.53 ± 0.04 | 82.8 |
| Exp3 | Z | 32 | 3 | 0 | 0 | – | – | – |
| Exp4 | T | 0.25 | 3 | 0 | 0 | – | – | – |
| Exp4 | T | 0.5 | 3 | 0 | 0 | – | – | – |
| Exp4 | T | 0.75 | 3 | 0 | 0 | – | – | – |
| Exp4 | Z | 0.25 | 2 | 2 | 1 | 116 (71–183) | 0.52 | 4.8 |
| Exp4 | Z | 0.5 | 2 | 2 | 2 | 103 (71–145) | 0.60 ± 0.08 | 7.4 |
| Exp4 | Z | 0.75 | 1 | 1 | 1 | 93 (56–146) | 0.64 | 8.5 |
| Exp4 | S | 0.25 | 3 | 2 | 2 | 91 (62–131) | 0.67 ± 0.07 | 7.6 |
| Exp4 | S | 0.5 | 3 | 3 | 1 | 98 (60–152) | 0.61 | 7.0 |
| Exp4 | S | 0.75 | 3 | 3 | 3 | 93 (59–142) | 0.67 ± 0.10 | 8.3 |
| Exp4 | S | 1 | 3 | 3 | 3 | 115 (85–155) | 0.52 ± 0.03 | 10.2 |
| Exp4 | S | 2 | 3 | 3 | 3 | 139 (115–167) | 0.45 ± 0.06 | 16.4 |
| Exp4 | S | 4 | 3 | 3 | 3 | 133 (113–155) | 0.45 ± 0.01 | 40.4 |

**Table ST3:** Fits for the pilot experiments (single wells). Reporter in µM; *τ* in min with 95% profile interval; rel.: *τ* passed quality control and was bounded.

| Exp | Src | U | Reporter | *τ* (min) | *k* (/h) | *A* | rel. |
|---|---|---|---|---|---|---|---|
| Exp1 | T | 2.5 | 5 | 46 (43–49) | 1.31 | 70 | yes |
| Exp1 | T | 2.5 | 2.5 | 44 (36–55) | 1.35 | 26 | yes |
| Exp1 | T | 2.5 | 1.25 | 20 (14–28) | 3.02 | 10 | yes |
| Exp1 | T | 2.5 | 0.83 | 13 (5–27) | 4.53 | 5 | no |
| Exp1 | T | 5 | 5 | 23 (21–27) | 2.58 | 101 | yes |
| Exp1 | T | 5 | 2.5 | 22 (20–24) | 2.69 | 45 | yes |
| Exp1 | T | 5 | 1.25 | 15 (12–18) | 4.04 | 18 | yes |
| Exp1 | T | 5 | 0.83 | 16 (11–23) | 3.71 | 9 | yes |
| Exp1 | T | 10 | 5 | 13 (10–17) | 4.54 | 124 | yes |
| Exp1 | T | 10 | 2.5 | 14 (11–18) | 4.25 | 61 | yes |
| Exp1 | T | 10 | 1.25 | 16 (13–21) | 3.69 | 27 | yes |
| Exp1 | T | 10 | 0.83 | 14 (11–19) | 4.21 | 30 | yes |
| Exp1 | W | 10 | 5 | 74 (66–84) | 0.81 | 94 | yes |
| Exp2 | G | 1.25 | 5 | 254 (186–359) | 0.24 | 7 | yes |
| Exp2 | G | 2.5 | 5 | 174 (144–209) | 0.35 | 18 | yes |
| Exp2 | G | 5 | 5 | 144 (132–157) | 0.42 | 75 | yes |
| Exp2 | T | 10 | 5 | 23 (18–27) | 2.66 | 169 | yes |
| Exp2 | W | 1.25 | 5 | 70 (61–80) | 0.86 | 139 | yes |
| Exp2 | W | 2.5 | 5 | 31 (25–37) | 1.96 | 145 | yes |
| Exp2 | W | 5 | 5 | 5 (2–7) | 10.99 | 113 | no |

**Figure SF1:**
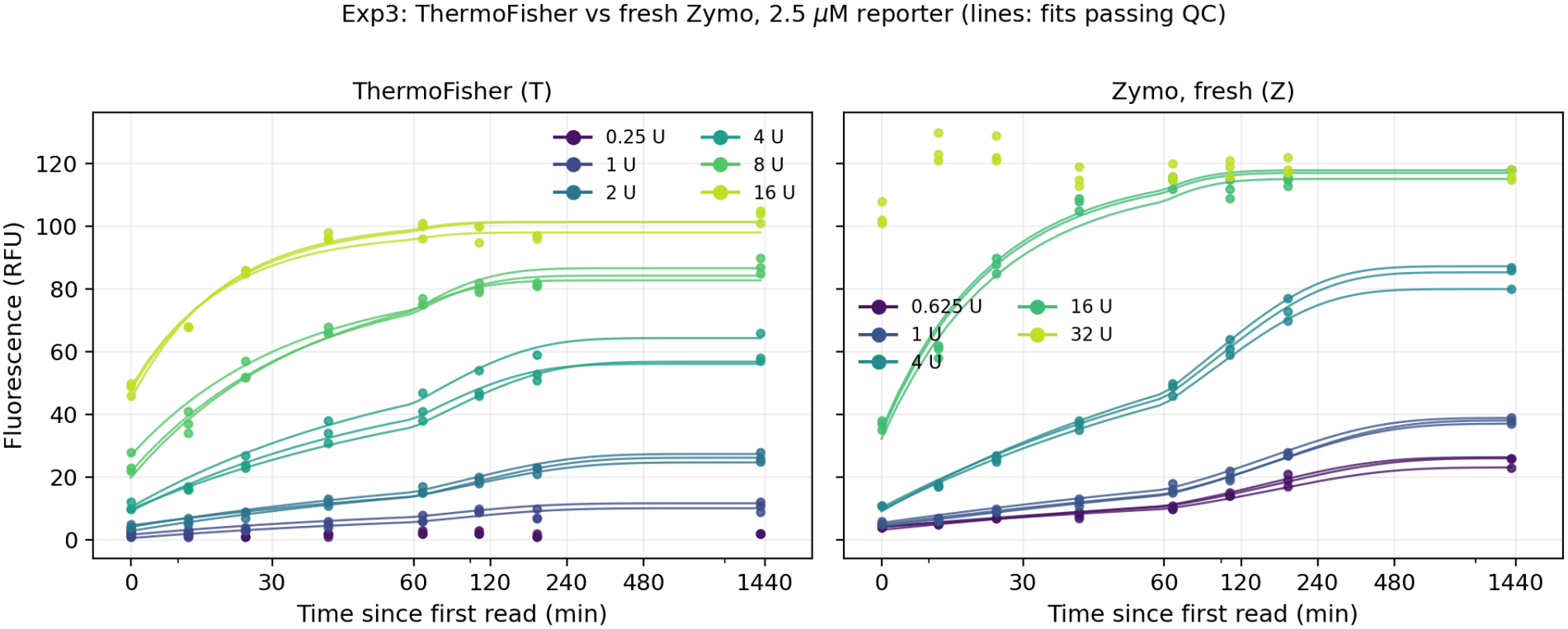
Exp3 time courses and fits. ThermoFisher and fresh Zymo, 2.5 µM reporter, all replicate wells, against recorded time (axis linear to 60 min, logarithmic beyond). Lines are fits passing quality control.

**Figure SF2:**
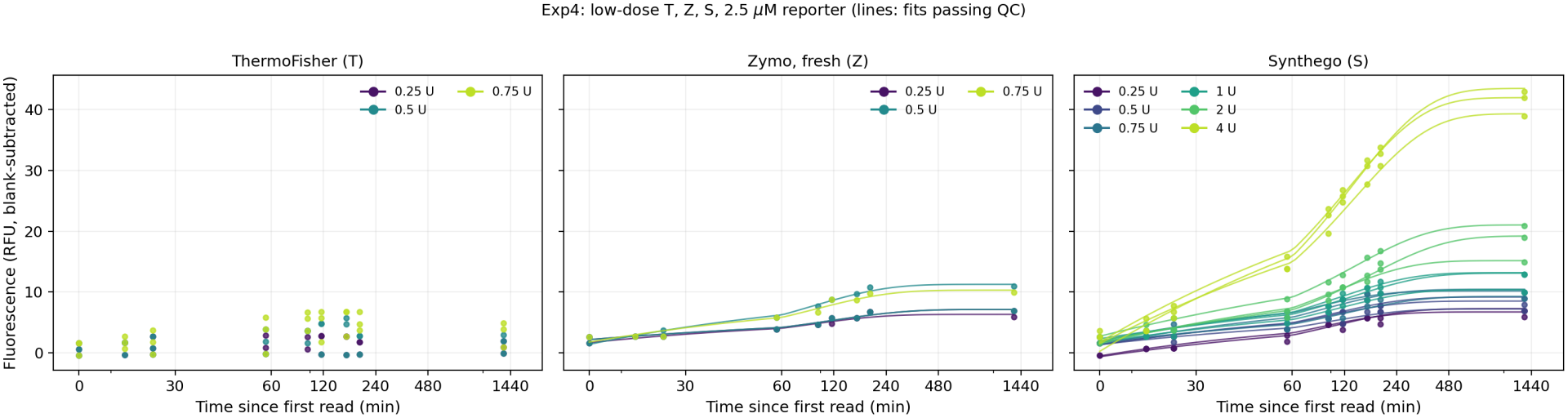
Exp4 time courses and fits. Low-amount ThermoFisher, Zymo and Synthego, 2.5 µM reporter, blank-subtracted. Most ThermoFisher wells at these amounts were indistinguishable from background and have no fit.

**Figure SF3:**
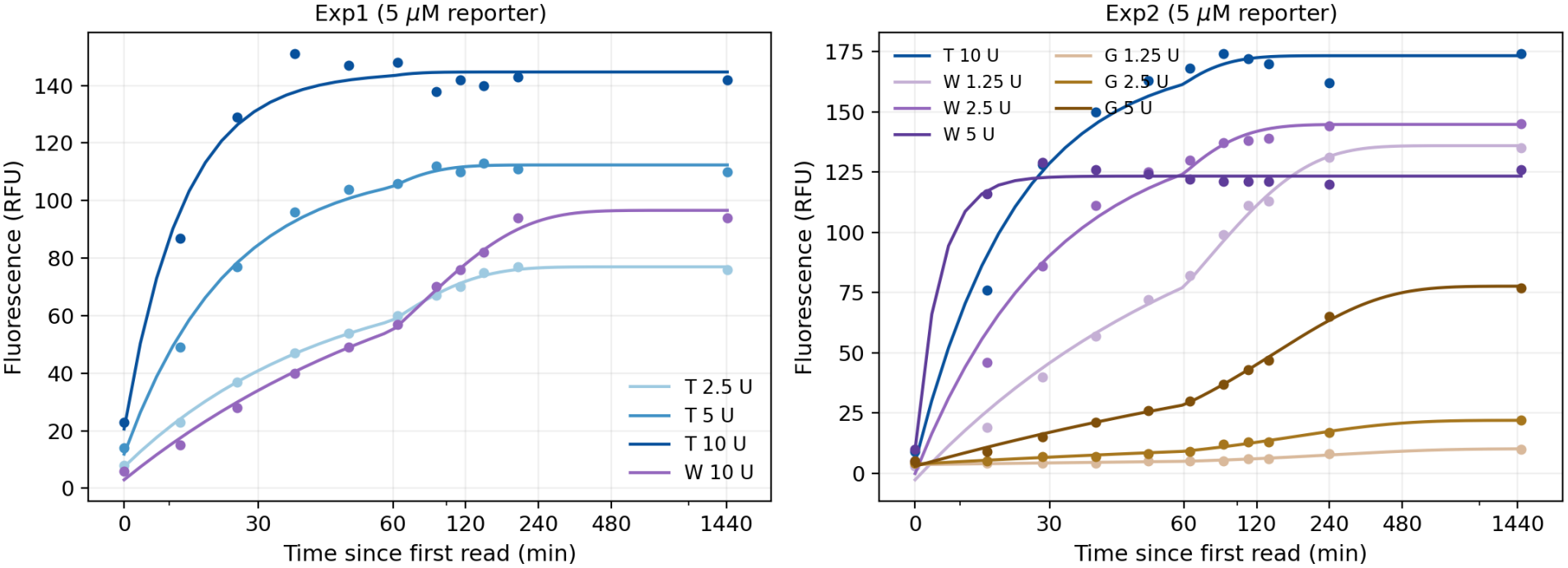
Pilot experiments. Exp1 (ThermoFisher at 2.5, 5 and 10 U and one W-aliquot well, 5 µM reporter) and Exp2 (ThermoFisher, and two Zymo aliquots W and G at 1.25–5 µL), single wells, against recorded time.

